# Ecological coupling of foraging and thermoregulation in harvester ants from a tropical deciduous forest in southern Mexico

**DOI:** 10.64898/2026.10.05.752865

**Authors:** Israel Valencia-Esquivel, Irma Inchaurregui, Anayelli Martínez, Dylan Zepeda, Elsah Arce, Marcela Osorio-Beristain

**Affiliations:** Centro de Investigación en Biodiversidad y Conservación, Universidad Autónoma del Estado de Morelos, Av. Universidad 1001, 62209, Cuernavaca, Morelos, México; Doctorado en Ciencias Naturales, Universidad Autónoma del Estado de Morelos, Cuernavaca, Morelos, México; Centro de Investigaciones Biológicas, Universidad Autónoma del Estado de Morelos, Río Pánuco 41, 62290, Cuernavaca, Morelos, México

**Keywords:** optimal foraging theory, thermal performance, habitat warming, climate change ecology

## Abstract

Diurnal ectotherms in hot and dry ecosystems face an acquisition trade-off between maximizing collective energy intake and minimizing individual exposure time. We employed optimal foraging and thermoregulation models to examine the surface foraging and thermoregulatory strategies of *Pogonomyrmex barbatus* foragers in response to habitat warming in a tropical deciduous forest. Foraging activity was evaluated using field-based thermal performance curves (fTPCs) for individual foraging speed and collective foraging success. Thermoregulation was assessed through indices of thermal unsuitability, as well as thermoregulatory effectiveness and exploitation. Our results show that afternoon conditions were significantly hotter and drier than those in the morning. Furthermore, the decoupling of fTPCs was dynamic; while the optimal temperature (T_opt_) for individual foraging speed remained stable at 44°C, the T_opt_ for collective foraging success shifted from 35°C in the morning to 41°C in the afternoon. This shift in foraging strategy, driven by stressful afternoon conditions, was reflected in a transition of the thermoregulatory strategy. Specifically, the probability of foraging activity decreased as thermal unsuitability increased, with a lower baseline in the afternoon. The thermoregulatory exploitation dropped from 94% in the morning to 63% in the afternoon. Based on thermoregulatory effectiveness, foragers transitioned from an active thermoregulatory strategy in the morning to a thermoconformist strategy in the afternoon while foraging on the surface. So, the observed colonies optimized the balance between individual foraging speed and collective foraging success through a thermoregulatory strategy that was primarily spatial in the morning and strictly temporal in the afternoon. A comparison with historical data from the same site reveals a 40% contraction in the total daily foraging window since 2013.

## Introduction

An ant colony functions as a superorganism organized into specialized castes (Mira et al. 2022), such as foragers, who are responsible for retrieving resources for the nest. This foraging behavior is linked to colony fitness and is thus under selection pressure. Optimal foraging models predict that selection should favor strategies that maximize the probability of foraging success (net energy intake) minimizing the associated cost (survival risk; Stephens & Krebs 1986). Foraging strategies of ants rely on behaviors and physiological processes that are thermodependent, so their performance—including cognition and movement— responds to the temperatures experienced by the ants (Huey 1991). Given this thermal dependence, models of optimal thermoregulation predict that natural selection promotes strategies that maximize ecological benefits (homeostasis, energy) minimizing the associated cost (survival risk; Angilletta 2009). Specifically, the extended cost-benefit model of thermoregulation predicts increasing thermoregulatory effort to maintain homeostasis by avoiding sublethal thermal conditions (Vickers et al. 2011). Thermoregulatory effort refers to behaviors such as seeking out cool areas while foraging on the ground, foraging during times when temperatures are suitable for doing so, or taking refuge inside the nest when experiencing thermal stress.

Foraging activity in social insects varies regarding multiple components, such as individual performance and collective coordination. At the individual level, foraging speed of foragers is largely dictated by temperature-dependent muscle contraction frequency (Hurlbert et al. 2008). This physiological trait allows *P. barbatus* foragers to reach their maximum foraging speed at a surface temperature of 45°C, close to that at which they lose motor coordination, i.e., their CTmax (50°C; Roeder et al. 2022). However, individual foraging speed does not translate linearly into foraging success at the colony level. Collective foraging success relies on a sensitive social feedback loop where the return rate of successful foragers stimulates the departure of new foragers (Schafer et al. 2006; Pinter-Wollman et al. 2013). Rising temperatures and declining humidity degrade this social feedback loop (Gordon et al. 2013; Friedman et al. 2019), possibly due to increase of overheating and desiccation risk (Menzel et al. 2026), and to the interference of chemical communication (Ruano et al. 1999). This cross-scale conflict should create a decoupling: while individuals possess the thermal tolerance to move rapidly in stressful hot conditions, the degradation of the social coordination should force the colony to restrict its activity to cooler microclimates. In tropical deciduous forests, the thermogenic constraint of the social coordination of *P. barbatus* foragers produces a bimodal activity pattern: foraging is concentrated in the morning (Ríos-Casanova et al. 2014), a window significantly cooler (−10.4°C) and more humid (+24.9%) than the afternoon (Hernández-Flores et al. 2016).

The thermal decoupling between foraging activity components and the circadian gradient of temperature and humidity generates two interconnected trade-offs. Under microclimates that represent a low hydrothermal stress, the physiological state of foraging ants (Menzel et al. 2026) allows them to maximize collective coordination (Gordon et al. 2013; Friedman et al. 2019) and communication (Ruano et al. 1999) during foraging, thus optimizing the probability of successful foraging without compromising the survival of workers. However, as temperature and dryness increase throughout the day, so does the risk of overheating and desiccation (Menzel et al. 2026). During these stressful conditions, foraging ants face an acquisition trade-off, continue foraging on the surface while experiencing sublethal conditions (Angilletta 2009). Regarding the extended cost-benefit model of thermoregulation, if the colony prioritizes current energy intake in stressful conditions, it risks foraging ants and future foraging capacity (Cunningham et al. 2021). Conversely, if foragers retreat to thermal refuges to ensure survival, they lose foraging opportunities (Cunningham et al. 2021). This shift leads to an allocation trade-off as ants must divide their time between food acquisition on the surface and thermoregulate inside the nest. So, the optimal temperatures for the colony during foraging must be the optimization of trade-offs between the individual foraging speed and collective foraging success.

In this study, we evaluated the foraging and thermoregulatory strategies involved in the acquisition and allocation trade-offs of the foraging activity of *P. barbatus* within a tropical deciduous forest. First, we analyze the simultaneous variation in individual foraging speed and collective foraging success throughout the temperature range during activity periods. Second, we analyze the thermoregulatory adjustments aimed at maintaining ideal temperatures for foraging activity during activity periods. According to the optimal foraging theory (Stephens & Krebs 1986) we expect that when conditions become thermally stressful, the optimal foraging strategy should shift from maximizing energy intake (consequence of maximizing collective foraging success) to minimizing individual exposure time (consequence of maximizing individual foraging speed) through increasing the thermoregulatory effort. As global warming continues to reshape habitat conditions, it is essential to understand how environmental shifts influence foraging performance of ants through thermoregulatory adjustments.

## Methods

### Study site

We conducted our study in April 2024, in the tropical deciduous forest within of the UNESCO Biosphere Reserve: Sierra de Huautla Biosphere Reserve. This reserve protects nearly 60,000 hectares of primarily tropical deciduous forest, where ectotherms find cooler conditions for thermoregulation compared to adjacent unprotected areas (Valencia-Esquivel et al. 2023), making it a climate regulator for the region. The tropical deciduous forest in the study region has two seasons, the hottest months occur from March to May, and the rainiest months occur from July to September (Valencia-Esquivel et al. 2023). The study region is characterized by a heterogeneous mosaic of preserved tropical deciduous forest, fragments under natural succession, areas where natural succession is arrested by livestock grazing, and land transformed to agriculture (Hernández-Flores et al. 2016). The study site region exhibits biodiversity of herbs, trees, and ants. Regarding the herbaceous stratum, a density of 3.6 (SD = 1.1) species/m² was recorded, of which three contributed most to the biomass: *Senna hirsuta, S. uniflora* (Fabaceae), and *Paspalum virgatum* (Poaceae). *Pogonomyrmex barbatus* ants forage seeds of *Senna* spp. and *Paspalum* spp. (de la O-Toris et al. 2012, Hernández-Flores et al. 2016). Regarding the tree stratum, a density of 252.5 trees/ha was recorded (Martinez-Garza et al. 2011). The most abundant tree species were *Vachellia campechiana, Ipomoea pauciflora, and Mimosa benthamii*. Regarding the ant community, 17 ant species were collected, of which *P. barbatus*, *Solenopsis geminata* (seed harvesters) and two *Pheidole* spp. (generalists) were the most abundant (Martinez-Garza et al. 2011).

### Experimental design

We recorded three equidistantly separated *P. barbatus* nests located near the El Limón Biological Station (18°32’31.18”N, 98°56’9.42”W, 1330 m a.s.l.). The open space surrounding the entrance of the ant nests was on average 6.6 m (SD= 1.3) long and 5.67 m (SD= 0.95) wide, respectively. The tree cover within a 10-meter radius of the nests was on average 28.5% (SD= 0.86). We recorded a total of 75 ant routes, with an average of 25 per nest (SD= 7.2). We proposed a design that simultaneously recorded individual foraging speed and collective foraging success, along with their associated temperatures, within the same spatiotemporal window. We recognize that this design limits inference at the population, inter-seasonal, and interannual levels, however, our design, was to measure the available thermal habitat for each individual worker during its foraging trip. First, we observed the nests’ entrance every 5 minutes between 8:30 h and 18:30 h to record the presence or absence of foraging activity to define the active foraging window. In the cleared area by the workers outside the nest, we look for foragers carrying an item during the activity periods. We recorded the duration time and distance traveled back to the nest to calculate their individual foraging speed (centimeters over seconds). We recorded the surface temperature every minute along the forager’s route with an infrared thermometer (emissivity set to 0.95). During the activity periods, we took a photograph every two minutes—in a radius of 10 cm from the nest entrance—to quantify collective foraging success as the proportion of loaded ants with items relative to the total foragers.

We characterized the available thermal habitat at different scales relevant to ants’ foraging activity. First, we estimated the thermohygrological conditions of the area surrounding the nests during observations. We calculated the surface temperature and relative humidity in the coordinates of the study area during the April days of 2024 using the micro_ncep function of the NicheMapR package (Kearney et al. 2020). We compare the surface temperature and relative humidity between morning and afternoon using a simple linear model. Second, during the ant’s activity periods, we recorded surface temperatures at the cells’ center of 5×5 m matrices centered on each nest. Third, we recorded surface temperatures of the nest entrances at 5-minute intervals throughout the day using autonomous thermometers (iButtons).

### Data analysis

#### Foraging strategy

We test the prediction that the optimal foraging strategy should shift from maximizing energy intake to minimizing individual exposure time when conditions become thermally stressful, as described below. To describe the foraging strategy, we estimated the field-based thermal performance curves of the collective foraging success as a proxy of energy intake and the individual foraging speed as a proxy of the individual exposure time. We determined the variation in individual foraging speed and collective foraging success during activity periods along the associated surface temperatures experienced by foragers in the field, i.e., their field-based thermal performance curves (fTPC; *sensu* Wild et al. 2025). We determined the fTPCs for individual foraging speed and collective foraging success using two quadratic models (following the recommendations in Angilletta 2006; Schwerdt et al. 2019; Tsai et al. 2020). First, we performed a mixed linear model on individual foraging speed relative to activity period (due to the reported bimodal activity pattern associated to the different environmental conditions), route temperature, and its quadratic term. Second, we performed a mixed logistic model on collective foraging success relative to activity period, nest entrance temperature, its quadratic term, and the interaction between activity period and nest entrance temperature. For all mixed models, we included nest identity as a random effect. We used the coefficients obtained from the quadratic models to calculate the optimal temperature (T_opt_) of the foraging activity components (following the recommendations in Schwerdt et al. 2019). The quadratic models describe a parabola, whose highest point (vertex) we defined as the T_opt_. We calculated the vertex as T_opt_ = −β1 / (2 * β2). We estimated the 95% CIs of the T_opt_ using the delta method of the car package in R. To test the prediction, we compared the distribution of fTPCs between the morning and the afternoon activity periods, this last when conditions should be hotter and drier. Specifically, we compared the distribution of temperatures along the routes used by foragers (for the individual foraging speed) and the temperatures of the matrix around the nests (for the collective foraging success) using the Kolmogorov-Smirnov test within each activity period. We used R to run the models (R Core Team 2020), and the following packages as performance for assessing whether the models meet the statistical assumptions (Lüdecke et al. 2021), car to get the confidence intervals (Fox & Weisberg 2019), ggplot2 to create the graphs (Wickham 2016), and emmeans to evaluate the post-hoc contrasts (Lenth 2024).

#### Thermoregulatory strategy

We test the prediction that optimal foraging strategy should consist in that investment in thermoregulatory effort should increase when conditions were thermally stressful during their foraging on the surface, as described below. We broke down the thermoregulation strategy into four indices: the (1) thermal activity breadth, (2) the probability of foraging activity related to the thermal unsuitability of the habitat, (3) thermoregulatory effectiveness, and (4) thermoregulatory exploitation index. First, we estimated the temperatures where the predicted probability of activity at the nest entrance was equal or higher than 50% calculating the thermal activity breadth of the foragers (TAB; *sensu* Cerdá et al. 1998). We considered the TAB as a proxy of the field-based thermal niche. To calculate the TAB, we performed a logistic mixed model on the activity status at the nest entrance relative to the surface temperature and its quadratic term. The model describes a parabola that intersects the 50% probability line of foraging activity at two consecutive points, which we considered as the lower and upper bounds of the TAB. To test the calculated upper TAB bound, we independently calculated the average of the maximum temperatures recorded for each of the foraging routes.

Second, according to our prediction, we determined whether in the afternoon nest entrance activity was more discouraged by the increase of the thermal unsuitability of the habitat than in the morning. We calculated the thermal unsuitability of the habitat, representing the coincidence of the available thermal habitat with the ants’ thermal niche, measured as the absolute deviation of matrix temperatures from the TAB. We fitted a logistic mixed model, with activity status at the nest entrance as the response, while thermal unsuitability of the habitat, activity period and their interaction were the fixed effects.

Third, thermoregulatory effectiveness was used to determine whether foragers deployed a thermoconformist behavior in the morning and a thermoregulatory behavior in afternoon while foraging on the surface, according to our prediction. To do so, we previously calculated the thermoregulatory inaccuracy of the ants (*following* Hertz et al. 1993). We calculated the thermoregulatory inaccuracy of the ants, representing the coincidence of the occupied thermal habitat with the ants’ thermal niche, measured as the absolute deviation of temperatures along foraging routes from the TAB. We then calculated thermoregulatory effectiveness as the thermal unsuitability of the habitat minus the thermoregulatory inaccuracy of the ants (*following the metric of* Nadeau & Blouin-Demers 2005). A thermoregulatory behavior was identified only when the thermoregulatory inaccuracy was significantly lower than the thermal unsuitability; otherwise, the lack of a significant difference indicated a thermoconformist behavior. For example, if the mean absolute deviation of available matrix temperatures from the TAB (e.g., 3°C) is significantly higher than the deviation of the occupied route temperatures (e.g., 2°C), we identify it as a thermoregulatory behavior (e.g., 1°C). To test our prediction with the thermoregulatory effectiveness, we constructed a generalized linear mixed model with a Tweedie distribution and a log link function to account for zero-inflated, continuous, positive absolute deviations. In this model, absolute thermal deviation was the response variable, while index type, activity period, and their interaction were fixed effects.

Fourth, thermoregulatory exploitation index (Ex; Taylor et al. 2021) was used to determine whether foragers were more efficient to exploit suitable foraging temperatures in the afternoon than in the morning. To do so, we filtered for the 5-minute intervals where the surface temperature at the nest entrance fell within the TAB (i.e., available opportunities). We then count the number of the intervals with (exploited opportunities) or without (unexploited opportunities) ants at the entrance of the nest, both in the morning and in the afternoon. Ex was analyzed as exploited opportunities over available opportunities, not as the proportion directly because multiple values can result in the same proportion. We fitted a binomial mixed model, with the exploited opportunities and available opportunities as the response and activity period as the fixed effect.

## Results

We identified two activity periods, a morning period (average duration of 1.64 h; 95% CI: 0.5–2.78) and an afternoon period (average duration of 1.22 h; 95% CI: 0–2.55; Figure 1). The afternoon presented hotter (+8.7°C; F= 55.68, p< 0.001; Table 1) and drier conditions (–25.2% relative humidity; F= 63.05, p< 0.001; Figure 1) than the morning. These microclimatic shifts corresponded with changes in the components of the foraging activity. The distribution of temperatures along the routes used by foragers and the temperatures of the matrix around the nests were decoupled in the morning (D= 0.33; p= 0.0003; Table 1), but coupled in the afternoon (D= 0; p= 1). In accordance, while individual foraging speed showed a stable thermal optimum of 43.9°C (95% CI: 41.2–46.5°C) between activity periods, collective foraging success varied with the interaction between activity period and nest entrance temperature (X^2^= 84.92, p< 0.001; Figure 2). Specifically, the optimal temperature for collective foraging success shifted from 34.96°C in the morning to 41.04°C in the afternoon, causing the intersection of the two field-based thermal performance curves (fTPCs) shifted from 38.1°C to 42°C (Figure 2). This represents a transition from thermally decoupled fTPCs in the morning—when speed operated at higher temperatures than success—to a convergence of performance curves in the hotter afternoon.

**Figure 1.**
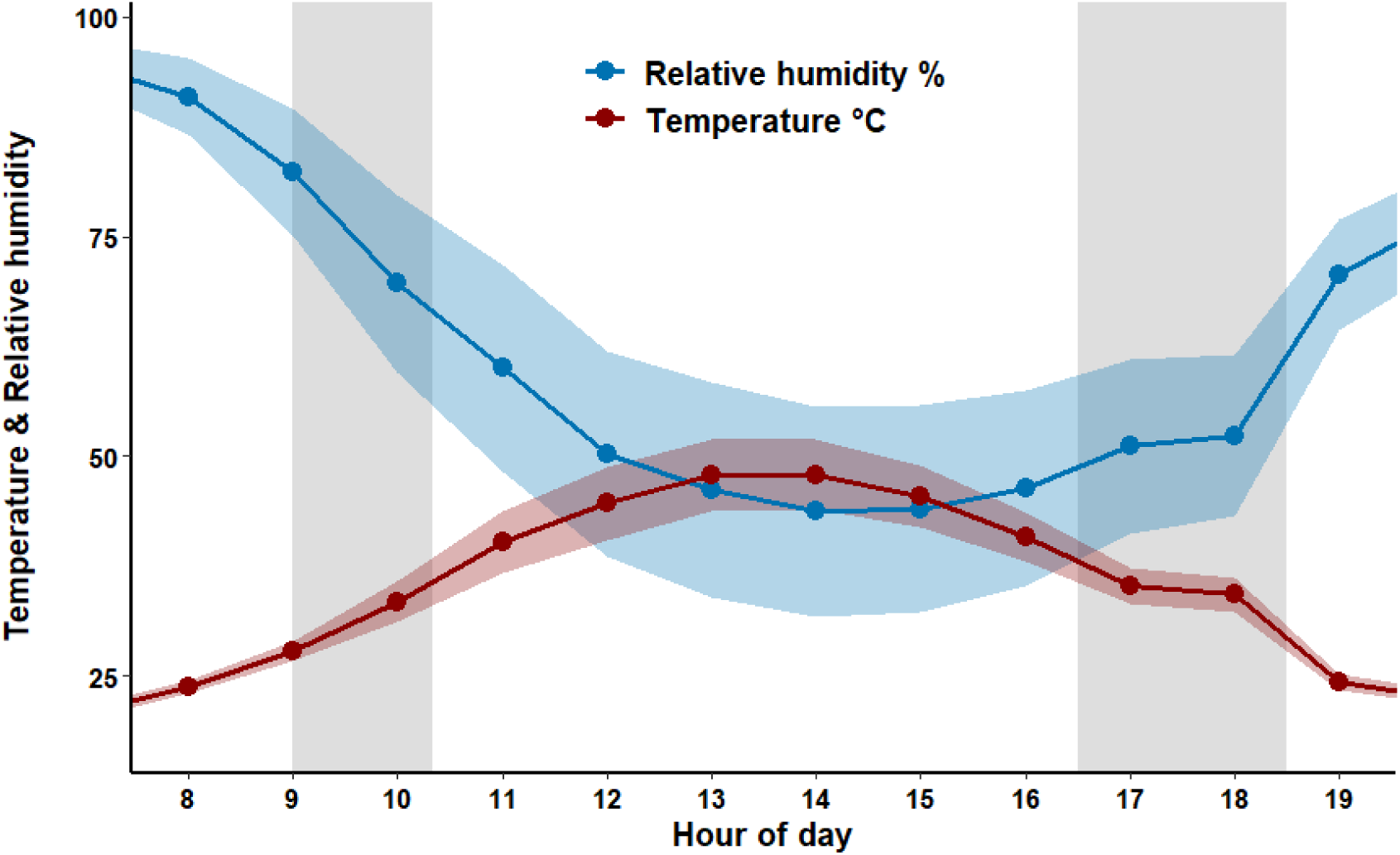
Climogram of average temperature and relative humidity during daylight hours. The areas around the lines correspond to their 95% CI. The gray vertical columns highlight the observed periods of morning and afternoon activity.

**Figure 2.**
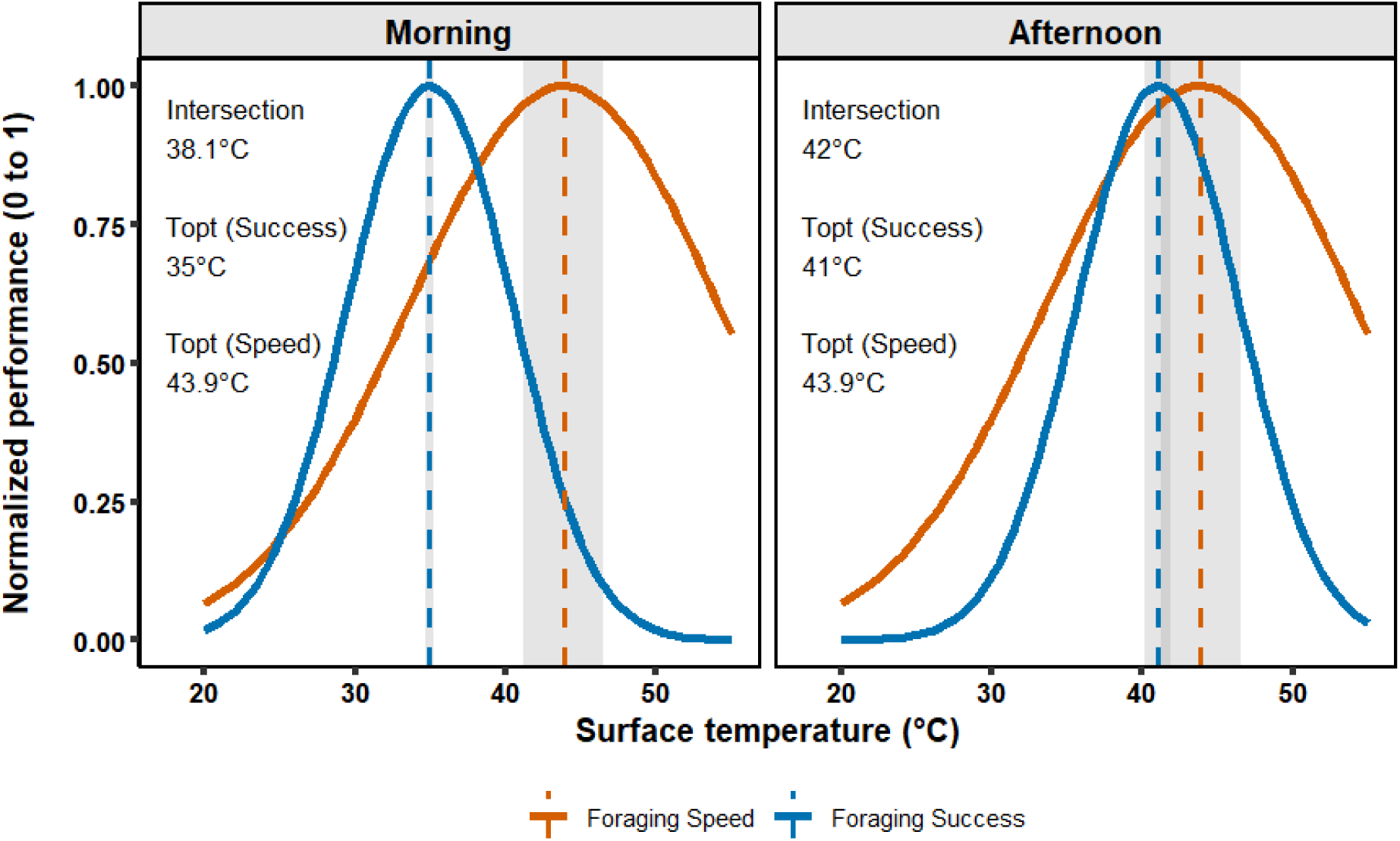
Field-based thermal performance curves (fTPCs) for collective foraging success (blue) and individual foraging speed (orange). The fTPCs represent the predicted values of the quadratic models described in Results. The optimal temperatures (Topt) are represented by vertical dotted lines corresponding to their color. The margins around the discontinuous lines represent their corresponding 95% CI. We standardized both fTPCs to a relative interval of 0 to 1, dividing the predicted values by their corresponding maximum.

**Table 1.**
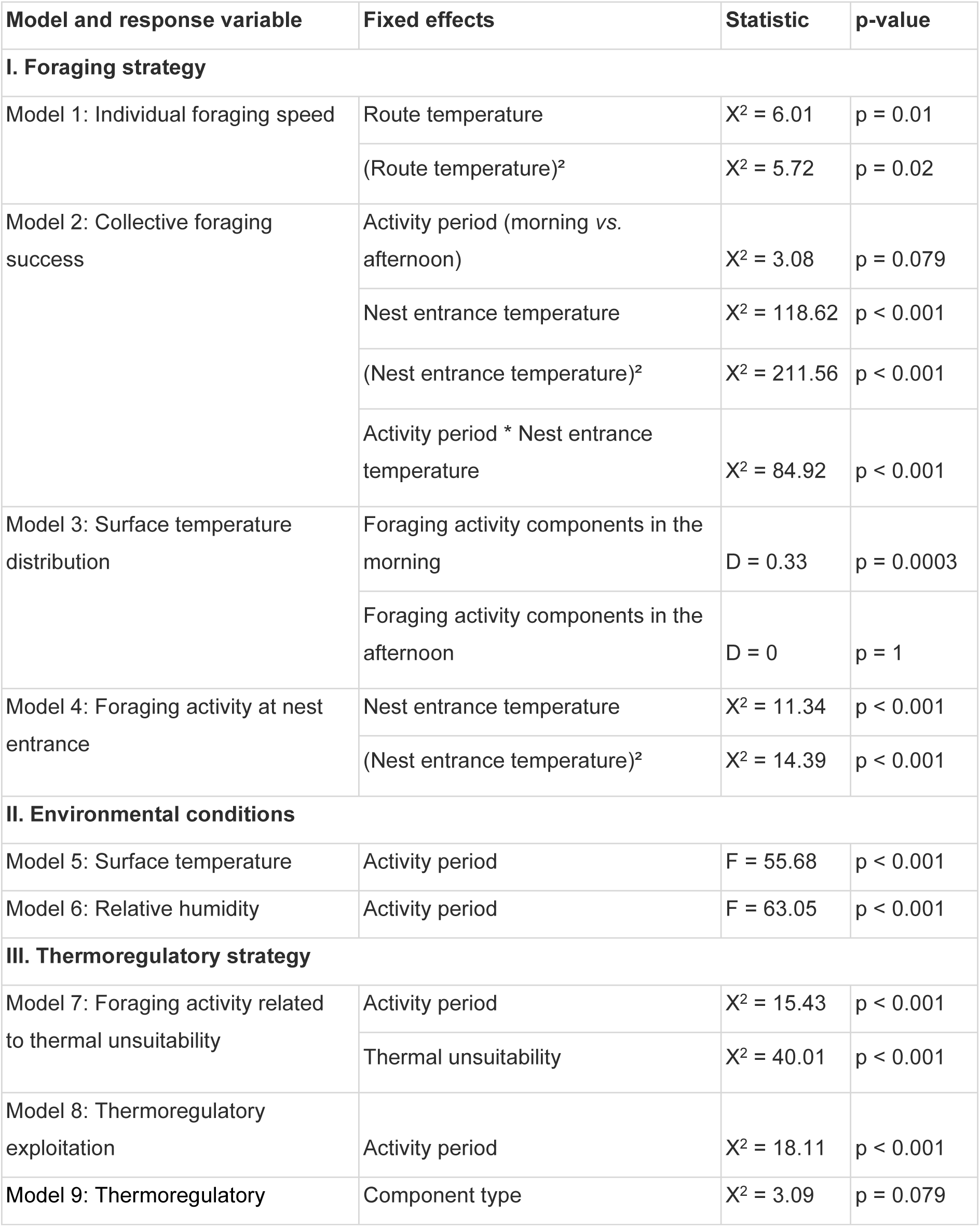
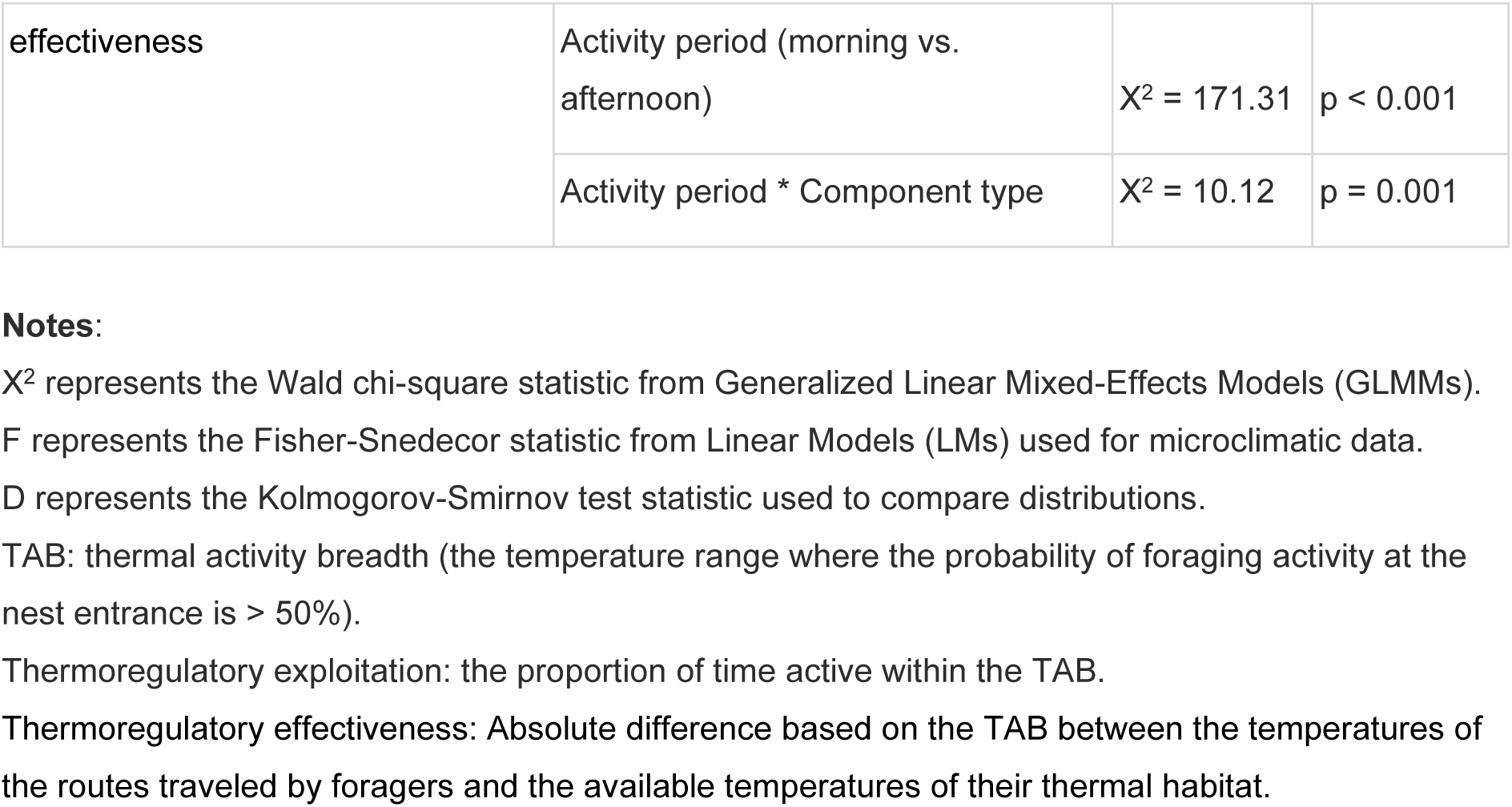
Summary of fixed effects for the linear and mixed-effects models evaluating the components of the foraging strategy and thermoregulatory strategy of *Pogonomyrmex barbatus* colonies, and environmental conditions.

Since the thermal activity breadth (TAB) indicates the temperatures at which ants are most likely (>50%) to forage, we use it to calculate the thermoregulation strategy indices. The TAB was bounded by 29.2°C and 43.7°C. The upper bound was independently mirrored by the maximum average temperatures recorded on foraging routes (43.74°C). The probability of the foraging activity decreased as the surface temperature at nest entrance deviated further from the TAB (X^2^= 40.01, p< 0.001; Table 1; Figure 3). The trend of the relationship was the same between morning and afternoon, but the intercept of the probability was lower in the afternoon than in the morning. Specifically, the probability of foraging activity dropped from 66% to 29% with a 1°C increase of the thermal unsuitability in the afternoon, compared to a drop from 89% to 64% in the morning (X^2^= 15.43, p< 0.001; Figure 3). This thermal restriction was not offset by a higher thermoregulatory effectiveness or thermoregulatory exploitation index (Ex). Thermoregulatory effectiveness—difference between habitat unsuitability and thermoregulatory inaccuracy—was significant in the morning (z= 3.07, p= 0.002; Table 1), but non-significant in the afternoon (z= −1.93, p= 0.054; Figure 4). Ex was 1.49 times higher in the morning (93.6%) than in the afternoon (62.7%; X^2^= 18.11, p< 0.001; Table 1). These indices suggest that *P. barbatus* colonies transition from a spatial thermoregulatory strategy during the morning to a temporal thermoregulatory strategy in the afternoon, when overheating and desiccation risks are higher.

**Figure 3.**
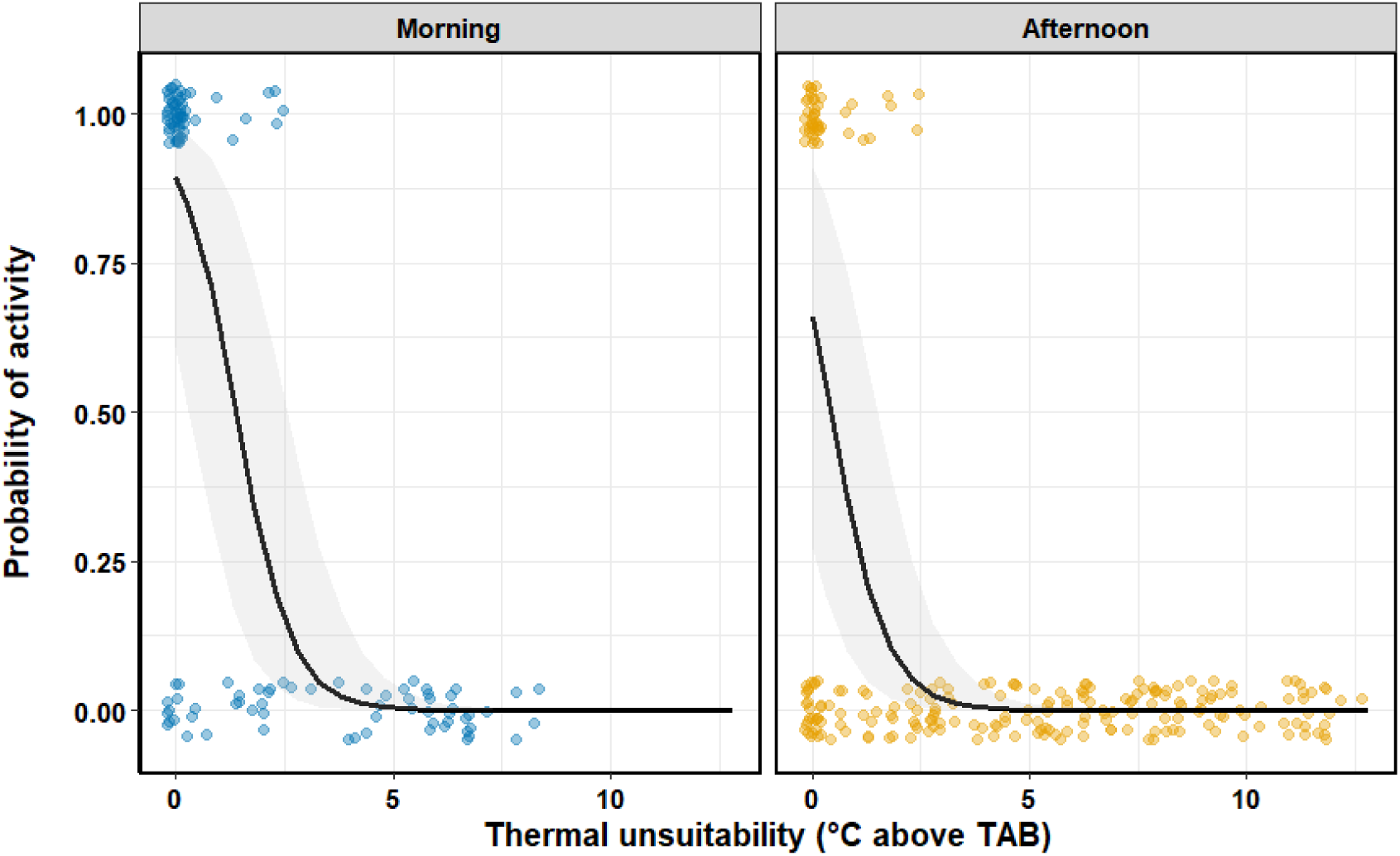
Probability of activity at the nest entrance as a function of thermal unsuitability during the morning and afternoon activity periods. The points represent observations of the presence (1) or absence (0) of ants at the nest entrance in the morning (blue points) and afternoon (yellow points). The black line indicates the model prediction, and the gray band indicates its 95% CI.

**Figure 4.**
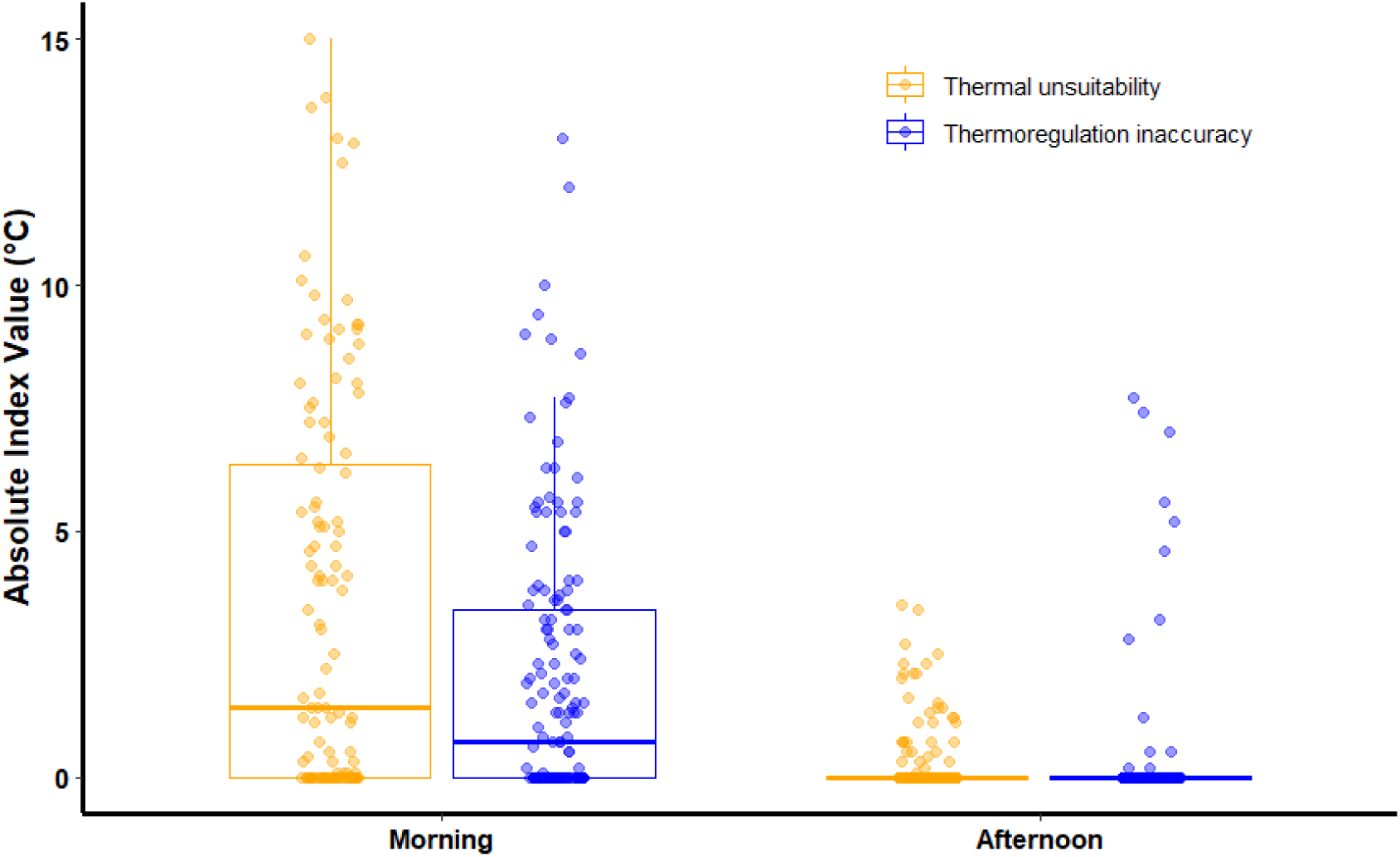
Thermoregulatory effectiveness during the morning and afternoon activity periods. The points represent observations of thermal unsuitability and thermoregulatory inaccuracy. The difference between both indices is the thermoregulatory effectiveness.

## Discussion

### Periods of foraging activity

The observed foragers had a bimodal activity pattern that was shorter in duration than that reported for other *Pogonomyrmex* colonies. The observed colonies presented a bimodal activity pattern, consistent with other *Pogonomyrmex* spp. (Pol & Casenave 2004; Ríos-Casanova 2014). While the threshold of the foraging activity pattern is traditionally attributed solely to desiccation tolerance (Menzel et al. 2025), our findings suggest it also reflects the optimization of the acquisition trade-off between a thermosensitive trait (collective foraging success) and a thermotolerant trait (individual foraging speed). Under this framework, selection favors the intersection of the field-based thermal performance curves (fTPCs), where the balance between individual foraging speed and collective foraging success is maximized. The temporal distribution of foraging activity on the surface of the observed ants should conform to this balance. On average, the observed ants foraged on the surface for 2.86 hours per day. This duration is almost three times shorter than the average reported in the literature for seven species of *Pogonomyrmex* (Table S1) in the dry season of arid ecosystems, 8.2 hours/day. The same pattern is repeated for five species of *Pogonomyrmex* when considering only the morning hours (Table S1); colonies of *Pogonomyrmex* spp. remain active for an average of 5.1 hours, tripling the duration of 1.64 hours observed in our colonies. The duration of foraging on the surface is even shorter than it was 11 years ago in the same area of this study. In 2013, *P. barbatus* at this location foraged for approximately six hours daily, from 08:00 to 11:00 h and 15:00 to 18:00 h (Hernández-Flores et al. 2016). 11 years later, the total daily foraging window has been reduced by more than 40%; the morning activity period has contracted—starting an hour later and ending 40 minutes earlier—while the afternoon emergence is now delayed by almost 90 minutes. Our data represents one of the shortest activity duration records documented for *Pogonomyrmex* due to experiencing the conditions of the driest month of the dry season tropical deciduous forest and its contraction over a decade.

### Thermal performance curves

As expected, when conditions became thermally stressful, the optimal foraging strategy shifted from maximizing energy intake to minimizing individual exposure time. The fTPCs of the foraging strategy changed between morning and afternoon conditions. The decoupling of the fTPCs of individual foraging speed and collective foraging success showed an acquisition trade-off during the morning. The intersection between both fTPCs occurred at 38.1°C, where the colony retained 80% performance in the individual foraging speed and collective foraging success. In contrast, the afternoon overlap of both fTPCs eliminated the acquisition trade-off. While the optimal temperature (T_opt_) of the individual foraging speed remained stable at 43.9°C, the T_opt_ of collective foraging success shifted upward to 41.04°C. The intersection of the acquisition trade-off did rise to 42°C. At this hotter threshold, the individuals must run at almost their maximum foraging speed to avoid the afternoon’s sublethal conditions while achieving the greatest possible collective foraging success under such conditions. This coincides with our prediction that as conditions become thermally stressful, the optimal foraging strategy should shift from maximizing energy intake to minimizing individual exposure time through increasing thermoregulatory effort. In another system, the foraging activity of *Myrmecia croslandi* and *M. pyriformis* ants decreases at temperatures lower than their foraging speed does in a dry sclerophyll forest (Jayatilaka et al. 2011). However, increasing temperature can both maximize energy intake and minimize individual exposure in ants. For example, in *Atta colombica* ants facing the cool, humid environments of the rainy season in a tropical rainforest (Welch et al. 2020). Even in *Dorymyrmex tener* ants facing experimental heating and feeding during the summer in a semi-arid temperate steppe (Devegili et al. 2025).

### Foraging activity based on thermal unsuitability

As predicted, thermoregulatory effort increased when conditions became thermally stressful regarding the thermal unsuitability and thermoregulatory exploitation. The probability of foraging activity decreased as the surface temperature deviated further from the thermal activity breadth. The baseline probability of foraging activity was lower in the afternoon than in the morning, therefore, for every °C that thermal unsuitability increased, the probability of foraging activity at the nest entrance was lower in the afternoon (−71%) than in the morning (−36%). Our results coincide with ant species in a subtropical grassland, which were less likely to occupy bait stations as the surface temperature deviated further from the species’ preferred temperature range (Youngsteadt et al. 2023). However, in the warmer adjacent urbanized environment, these ants did not occupy the available cool sites or times to mitigate the hotter conditions. In our study, foragers did not compensate for the lower probability of foraging success in the afternoon with a greater thermoregulatory exploitation (i.e., a higher proportion of activity during the time with conditions suitable for foraging) or thermoregulatory effectiveness (i.e., a deviation of the temperatures occupied by ants from the TAB lower than expected).

### Thermoregulatory effectiveness

As we predicted, thermoregulatory effort was higher in the morning, than in the afternoon, regarding the thermoregulatory effectiveness index. During the afternoon, the synergy between hot and dry conditions likely increased the desiccation rate of foragers (Friedman et al. 2019), reducing their thermal tolerance. Under these conditions, the water balance of foraging becomes less efficient (Friedman et al. 2019). As a result, foragers emerged from the nest to forage in the afternoon only when route temperatures remained as close to the thermal activity breadth as the available habitat temperatures. According to the thermoregulatory effectiveness, this suggests a thermoconformist behavior; however, in fact, foragers exhibited a temporal, rather than spatial, thermoregulatory strategy during the afternoon. Conversely, the cool, humid morning conditions allowed foragers to occupy routes cooler than the available thermal habitat, thereby expanding their foraging window even when matrix temperatures exceeded the thermal activity breadth. This explains why thermal unsuitability was greater in the morning than in the afternoon. Given the suitable conditions of the morning, we suggest that our colonies could afford to minimize foraging during the afternoon because the high profitability of granivory in *Pogonomyrmex* spp. (Weier & Feener 1995) allowed them to satisfy most of their daily energy requirements during the morning. The optimal foraging strategy for our colonies was therefore to distribute foraging activity based on the risk of desiccation and overheating during exposure.

Our results allow us to contrast two classic models about the behavioral response of ectotherms to stressful temperatures, the classic (Huey & Slatkin 1976) and extended (Vickers et al. 2011) cost-benefit model of thermoregulation. Our results support the extended cost-benefit models of thermoregulation (Vickers et al. 2011), which predicts that thermoregulatory effort increases to avoid thermogenic effects. Accordingly, foragers deployed a thermoregulatory strategy that was primarily spatial in the morning, but transitioned to a strictly temporal one in the afternoon. Such afternoon strategy implies losing foraging opportunities, leading to an allocation trade-off between time spent foraging on the surface and the time spent sheltering inside the nest (Cunningham et al. 2021). Nevertheless, the foragers did not compensate for losing foraging opportunities in the afternoon by making more efficient use of periods with temperatures conducive to foraging (Ex). Ex indicates selection pressures, unrelated to habitat temperature, that limit the ability to achieve a thermoregulatory strategy (Taylor et al. 2021). Undoubtedly, the dry conditions in the afternoon reduced heat tolerance and increased the risk of experiencing sublethal conditions, which prevented the proposed compensation. This is concerning because understanding the effect of climate change on ant-mediated ecological processes—such as seed dispersal, arthropod predation, and soil aeration—requires the understanding their thermal sensitivity (*sensu* Williams et al. 2008). That is, the thermal dependence of their ecological processes and their associated thermoregulatory behaviors. Apart from the study by Youngsteadt et al. (2023), and to the best of our knowledge, there are no other studies against which we can compare the thermoregulatory strategy of ants while foraging on the ground—using the current thermoregulatory indexes.

### Limitations of this study

The design allowed us to simultaneously record, within the same spatial-temporal framework, individual foraging speed and collective foraging success along with their associated temperatures: the temperature along the routes, the open space around the nest, and the nest entrance. This high spatial and temporal resolution was necessary to link microclimatic conditions to the acquisition and allocation trade-offs of the foraging activity. However, we recognize that the strengths of this mechanistic approach also limit inference at the population level. There may be inter-colony variability in the behavioral variables evaluated, associated with factors such as colony size, nutritional status, or local thermal history (*see* Parr & Bishop 2022). We conducted the study during the dry season, so we do not know whether the acquisition trade-off between individual foraging speed and collective foraging success persists during the rainy season, when lower temperatures and higher humidity could reduce the risk of overheating and dehydration (*see* Welch et al. 2020). The interannual consistency of our results requires long-term validation given intraannual phenology and interannual climate variability (Gordon et al. 2013). Future studies can evaluate the dynamic of the observed acquisition trade-off and its associated foraging and thermoregulatory strategies through a larger number of colonies and on an inter-seasonal and intra-annual basis.

### Implications of global warming

Thermoregulation strategies associated with observed foraging strategies place the studied colonies in a complex position of vulnerability to climate warming. Under a time-for-time substitution model (*sensu* Lovell et al. 2023), we project that warming will cause the mild conditions of the morning activity period to approach the tolerable conditions of the current afternoon activity period, which in turn will be as thermally unviable as the current midday inactivity period. This loss of foraging activity time could be compensated in two ways. First, through the current temporal and spatial thermoregulation strategies deployed by the observed foraging ants. Second, expanding activity hours to cooler and more humid times of day. Although there are records of *P. barbatus* (Whitford et al. 1976) and *P. rugosus* (Mehlhop et al. 1983) foraging after dusk— taking advantage of the thermal inertia of the soil and the recovery of moisture—we are not aware of any reports of activity before dawn. We therefore predict an asymmetric response: the afternoon period could shift toward nighttime, but morning activity would be forced to concentrate in its current window without expanding towards dawn.

## Supplementary material

**Table S1.** Foraging duration in different *Pogonomyrmex* species inhabiting desert ecosystems or dry forests during the dry season. The records are sorted by the time of day when the ant nests were observed, either throughout the daylight hours (diurnal) or only in the morning. The records are then listed alphabetically by species.

| Species | Foraging duration (hours) | Part of the day | Citation |
| --- | --- | --- | --- |
| *P. barbatus* | 9 | Diurnal | Ríos-casanova 2014 |
| *P. barbatus* | 7 | Diurnal | Whitford et al. 1976 |
| *P. mayri* | 11 | Diurnal | Kugler 1984 |
| *P. naegelii* | 7 | Diurnal | Belchior et al. 2012 |
| *P. occidentalis* | 6 | Diurnal | Cristl et al. 1991 |
| *P. pronotalis* | 9.5 | Diurnal | Pol et al. 2004 |
| *P. rastratus* | 9 | Diurnal | Pol et al. 2004 |
| *P. rugosus* | 7 | Diurnal | Whitford et al. 1976 |
| P. barbatus | 6 | Morning | Gordon 1986 |
| P. barbatus | 6 | Morning | Gordon 1984 |
| P. barbatus | 5 | Morning | Gordon 1983 |
| P. californicus | 5 | Morning | Gordon 1984 |
| P. desertorum | 5 | Morning | Gordon 1984 |
| P. maricopa | 5 | Morning | Gordon 1984 |
| P. rugosus | 4 | Morning | Gordon 1984 |

## Notes

### Competing Interest Statement

The authors have declared no competing interest.

